# Predicting depression risk in early adolescence via multimodal brain imaging

**DOI:** 10.1101/2023.04.10.536286

**Authors:** Zeus Gracia-Tabuenca, Elise B. Barbeau, Yu Xia, Xiaoqian Chai

**Affiliations:** Department of Statistical Methods, University of Zaragoza, Zaragoza, Spain; Department of Neurology and Neurosurgery, McGill University, Montreal, Quebec, Canada; Department of Bioengineering, McGill University, Montreal, Quebec, Canada

## Abstract

Depression is an incapacitating psychiatric disorder with high prevalence in adolescent populations that is influenced by many risk factors, including family history of depression. The ability to predict who may develop depression before adolescence, when rates of depression increase markedly, is important for early intervention and prevention. Using a large longitudinal sample from the Adolescent Brain Cognitive Development (ABCD) Study (2658 participants after imaging quality control, between 9-10 years at baseline), we applied machine learning methods on a set of comprehensive multimodal neuroimaging features to predict depression risk at the two-year follow-up from the baseline visit. Features include derivatives from structural MRI, diffusion tensor imaging, and task and rest functional MRI. A rigorous cross-validation method of leave-one-site-out was used. Additionally, we tested the prediction models in a high-risk group of participants with parental history of depression (N=625). The results showed all brain features had prediction scores significantly better than expected by chance. When predicting depression onset in the high-risk group, brain features from resting-state functional connectomes showed the best classification performance, outperforming other brain features based on structural MRI and task-based fMRI. Results demonstrate that the functional connectivity of the brain can predict the risk of depression in early adolescence better than other univariate neuroimaging derivatives, highlighting the key role of the interacting elements of the connectome capturing more individual variability in psychopathology compared to measures of single brain regions.

## 1. INTRODUCTION

Adolescence is a period in life with substantial neural and hormonal changes which are also accompanied with noticeable changes in behavior. However, some of these changes may result in maladaptive or unpleasant behaviors that can lead to long-term effects with its consequent negative impact on individuals and their families and communities. In fact, many psychiatric disorders have their onset in the adolescent period (Kessler et al., 2005; Paus et al., 2008). Given that, it is of great importance for biomedical sciences to predict the risk for psychiatric disease onset in adolescents. Achieving that, more effective preventions and treatments can be applied, with consequent improvement in the quality of life for those individuals at risk.

Depression is a prevalent psychiatric disorder, highly recurrent and with a negative impact on quality of life, and can lead to mortality (Kessler et al., 2013; Tolentino and Schmidt, 2018). It has a broad spectrum of symptoms that may include anhedonia, sleep difficulties, and suicidal ideation, among others (Tolentino and Schmidt, 2018), and is considered the most prevalent cause of disability worldwide (Smith, 2013; Friedrich, 2017). Many variables contribute to a higher risk of developing depression including genetics, socio-economic factors, or environmental stress (Hammen, 2018; Zajkowska et al., 2021). Among these risk factors, parental history of depression significantly increases the risk of depression in offspring to three to five folds higher than individuals without a parental history of depression (Lieb et al., 2002; Weissman et al., 2016). Therefore, it is especially important to identify biomarkers for depression in this high-risk population of adolescents with parental history of depression. From a human neuroscience perspective, several studies have focused on the prediction of depression based on neuroimaging data (MacQueen, 2009; Nouretdinov et al., 2011; Gao et al., 2018). However, most studies focused on adult samples and how they differ from (well-balanced) control participants. In addition, these studies were limited by relatively small sample sizes and/or, importantly, the lack of longitudinal designs. Longitudinal designs are a fundamental tool to describe prospective lifespan changes at the individual level (Chen et al., 2021; Gracia-Tabuenca et al., 2021). Considering the importance of predicting the onset of depression, more research is needed in early or pre-adolescent groups from large community-based samples which better represent the population.

These limitations can be overcome with recent large scale open science projects. Particularly, to address the better characterization of the onset of psychiatric disorder in the early stages of adolescence, the ABCD Study encompasses longitudinal follow-ups of ∼12k participants of 9-10 years old accounting for their phenotypical, neuroimaging, and genetic assessment (Volkow et al., 2018). Two recent studies have evaluated the onset of depression in open datasets in adolescent samples using neuroimaging features, with modest prediction performances (Toenders et al., 2019, 2022; Ho et al., 2022). One study combined structural Magnetic Resonance Imaging (MRI), clinical and environment assessment scores in 14-year-old adolescents from the IMAGEN consortium data to predict the onset of major depression at 2- and 5-year follow-up (Toenders et al., 2022). They reported that baseline depression severity at age 14, female sex, neuroticism, stressful life events, and surface area of the supramarginal gyrus were the strongest predictors for depression onset. However, functional MRI features were not included as predictors. A second study by Ho et al. examined depression symptoms from a younger sample of 9- and 10-year-old children from the ABCD study and found that parental mental health, family environment, and child sleep quality were the top predictors of depression symptoms at the baseline and at the 1-year follow-up, while brain features had relatively weaker prediction power (resting-state functional MRI) or little to no predictive power (structural MRI) (Ho et al., 2022). These studies identified possible behavioral, demographic and environmental risk factors for depression and provided early evidence for different brain imaging markers for depression in adolescents.

The present study aims to predict the onset of depression in early adolescence based on a set of comprehensive brain features measured by multimodal MRI. To do so, we included the ABCD study data from the baseline visit and 2-year follow-up, when a larger proportion of participants have developed depression, compared to the 1-year follow-up investigated by Ho et al. (2022). We applied multivariate techniques to extract features from structural, diffusion-weighted, and (rest and task) functional MRI at the baseline visit and tested how well these different types of brain features predict depression onset at the two-year follow-up. We are particularly interested in vulnerability factors in the subsample of participants with parental history of depression, given the higher risk of developing depression in this subsample of children (Lieb et al., 2002; Weissman .et al., 2016; Ho et al., 2022). In sum, this study uses the biggest longitudinal early adolescent neuroimaging sample currently available to predict the onset of depression at the two-year follow-up, by integrating and comparing a comprehensive set of multimodal MRI brain imaging features, using a rigorous cross-validation method (leave-one-site-out) and focusing on a high-risk group of adolescents with familial risk of depression.

### 1. 2. METHODS

#### 2.1 Sample

The sample of the ABCD Study includes 11875 participants between 9 and 10 years old at baseline who are followed periodically in an intended span of 10 years. Baseline sampling occurred between September 2016 and August 2018 through 21 sites distributed across the United States of America. The study was approved by each site’s Institutional Review Board (Garavan et al., 2018). The final set of participants included in this study are described in section 2.8 below.

For this study, phenotypic data at baseline and at the two-year follow-up was extracted from the ABCD 3.0 release from the National Institute of Mental Health Data Archive (NDA) repository, as well as the baseline derivatives available from the MRI data: structural MRI, diffusion-weighted imaging, and task-based and rest fMRI. Additionally, preprocessed functional connectivity matrices at baseline were extracted from the ABCD-BIDS Community Collection (ABCD collection 3165; https://github.com/ABCD-STUDY/nda-abcd-collection-3165).

#### 2.2 MRI acquisition

Imaging protocol was harmonized for three types of 3 Tesla MR manufacturers (General Electric, Phillips, and Siemens). T1- and T2-weighted sequences were 1 mm3 isometric size. Diffusion-weighted images (DWI) were 1.7 mm3 isometric size with multi-band acceleration factor of 3, and 96 directions at different b-values (0, 500, 1000, 2000, and 3000). Functional MR images (fMRI) consisted in gradient-echo EPI (Echo Planar Images) with 2.4 mm3 isometric size, multi-band acceleration factor of 6, repetition time TR=800 ms, and echo time TE=30 ms. More information about MR sequences can be found at Casey et al. (2018).

#### 2.3 fMRI paradigms

fMRI sequences include four 5-minute runs of resting condition and three task-specific paradigms with two runs each. In the rest runs participants were asked to remain still with their eyes open while seeing a fixation crosshair. Task fMRI paradigms include a Monetary Incentive Delay (MID) task, a stop signal task (SST), and an emotional N-back task. For detailed descriptions of the tasks, please see supplementary methods material.

#### 2.4 MRI preprocessing

More detailed information about MRI preprocessing can be found at Hagler et al. (2019). Briefly, structural MRI (sMRI) underwent scanner-specific gradient nonlinearity distortion correction, intensity inhomogeneity correction via B1-bias field estimation, and were registered and resampled to an isotropic standard space. Cortical surface and subcortical segmentations were extracted using FreeSurfer v5.3. Structural measurements include cortical thickness, area, volume, sulcal depth, and intensity for T1w, T2w, and T1w/T2w ratio. Also, weighted averages for fuzzy-cluster parcellations based genetic correlation were computed (Chen et al., 2012), as well as intensity scores for the volumetric subcortical regions were included.

DWI were corrected for eddy currents, motion, susceptibility distortion, and gradient nonlinearity distortions. Major white matter fiber tracts were segmented using AtlasTrack. Diffusion Tensor Imaging (DTI) analysis were applied and standard measures were extracted: fractional anisotropy (FA), mean (MD), longitudinal (LD), and transverse (TD) diffusivity.

fMRI volumes were corrected for motion, susceptibility distortion, and gradient nonlinearity distortions. Task-fMRI contrasts were assessed using a general linear model (GLM) and calculated for each region of interest (ROI). GLM included as covariates baseline, quadratic trends, motion estimates, and their derivatives. Motion covariables were band-pass filtered at 0.31–0.43 Hz using an infinite impulse response (IIR) filter. Also, time points with a framewise displacement (FD) above 0.9 mm were censored. Rest-fMRI underwent additional preprocessing steps including removal of initial volumes, normalization, regression, temporal filtering (0.009 - 0.08 Hz), and calculation of average ROI time series. Functional connectivity matrices were extracted via Pearson’s cross-correlation of the Gordon’s parcellation ROIs. Time points with FD higher to 0.2mm were not included in the correlation. Fisher’s transform was applied to the correlations and were averaged based on the fourteen resting-state network predefined modules.

In addition to network-level resting-state connectivity matrix described above, we also extracted the ROI-to-ROI rest-fMRI connectivity matrices that were preprocessed following the ABCD-BIDS pipelines (https://github.com/DCAN-Labs/abcd-hcp-pipeline; DOI: https://doi.org/10.5281/zenodo.5757650). From a subset of 10073 scans from baseline, those with at least 8 minutes of uncensored time series were selected (N = 5955). These are Fisher’s r-to-z transformed functional connectivity matrices of pairwise connections between 352 regions of interests (ROIs). These ROIs include the 333 areas from Gordon et al. (2016) plus 19 subcortical regions from the FreeSurfer segmentation.

#### 2.5 Target variable

The target variable for the prediction models is the depression onset, defined as a binary variable if any of the 8 diagnostic scores of the parental Kiddie Schedule for Affective Disorders and Schizophrenia (KSADS-5; Kaufman et al., 2013) was positive at the two-year follow-up. The scores included persistent depressive disorder past, present or in partial remission; major depression disorder past, present or current in partial remission; and unspecified depressive disorder past or current. At two-year follow-up, data from 6317 matched participants was available (depression onset: Yes = 358; No = 5959; prevalence = 5.67%).

#### 2.6 High-risk group with parental history of depression

Considering previous findings of higher vulnerability for depression risk in subjects with parental history of depression (Lieb et al., 2002; Weissman et al., 2016), we further tested a subset of participants who reported maternal and/or paternal history of depression based on parental responses on the ABCD Family History Assessment. From the previous sample, a subset of 1854 participants (depression onset: Yes = 195; No = 1656; prevalence = 10.52%) from the two-year follow-up had parental history of depression.

#### 2.7 Predictors

All predictor variables were selected from baseline, this way the classification tests the prospective prediction of depression onset at the two-year follow-up. Brain features include: 1196 from sMRI, 1140 from DTI, 2548 from task-fMRI, and 61776 from rest-fMRI.

sMRI features included structural measures of 71 cortical thickness regions from the Desikan-Killiany atlas (Desikan et al., 2006) plus 36 weighted average thickness for the genetically derived fuzzy-cluster parcellations (2, 4, and 12 clusters) (Chen et al., 2012). For these 107 regions, four features were selected: cortical thickness, sulcal depth, surface area, gray matter volume. Additionally, for these 107 ROIs, another six features for (three for each T1 and T2 images) were extracted from the average intensity of white matter (voxels 0.2 mm from the white matter surface), gray matter (voxels 0.2 mm from the white matter surface), and white-gray contrast. Finally, from the 36 subcortical regions, three features were selected: volume, and T1 and T2 average intensity.

DTI features included four standard measures (FA: fractional anisotropy; MD: mean diffusivity; LD: longitudinal diffusivity; and TD: transverse diffusivity) extracted from 42 tracts and 30 subcortical regions, plus the sub/adjacent white-matter, cortical gray matter, and gray/white matter contrast associated with 71 cortical regions (i.e., three times 71 cortical features) resulting in a total of 1140 features. Major white fiber tracts were labeled via AtlasTrack (Hagler et al., 2009), while cortical and subcortical ROIs were extracted from the Desikan-Killiany atlas (Desikan et al., 2006).

Task-fMRI features consisted of the two-run average beta weight divided by its standard error from its corresponding contrast within 68 cortical plus 30 subcortical regions from the Desikan-Killiany atlas (Desikan et al., 2006). Specifically, MID, SST, and N-Back task-fMRI include 980, 686, and 882 features resulting from their corresponding ten, seven, and nine contrasts, respectively. The resulting 2548 variables were pooled together for the prediction analysis.

Rest-fMRI features were derived from the 61776 pairwise functional connectivity variables from the upper triangle of the preprocessed 352×352 ROI-ROI connectivity matrix. In addition, those ROI pairs were summarized into 13 resting-state networks (RSN) according to Gordon et al. (2016), then a total of 91 possible RSN interactions were selected (including also within-RSN summarized connectivity). The summarized network-level connectivity values were extracted from the tabulated ABCD baseline data.

#### 2.8 Inclusion/exclusion criteria

From the full sample at baseline, 6392 participants with at least 8 minutes of non-motion affected rest-fMRI data were selected. To avoid potential confounding effects of the bipolar disorder, 929 participants were excluded due to positive scores in the diagnostic bipolar variable in either the parental or youth report on the K-SADS. Additionally, 177 participants with conflicting scores between the child and parental report were excluded (a positive score in the depression onset variable from the youth’s KSADS responses when their parental responses were negative). From those, 3085 had available corresponding target data (KSADS-5) at two-year follow-up. 159 participants with depression at baseline were removed. Furthermore, 128 participants were removed due to missing data in any of the features. Finally, sites without positive cases were excluded for the prediction analyses, that is three sites with a total 140 participants.

The final dataset used in the classification analyses included a general sample of 2658 participants from 18 sites with complete data at baseline and at the two-year follow-up. From these participants, the high-risk group of parental history of depression includes 625 participants. Table 1 summarizes the sample selection.

**Table 1.** Number of participants with positive/negative depression based on the KSADS scores, total number of participants, and depression prevalence, for the general group as well as the high-risk group (with parental history of depression). A) Available sample at baseline with complete KSADS; B) Baseline sample matched with quality controlled (QC) MRI data; C) Final sample with complete data at baseline and two-year follow-up, no depression at baseline and negative scores in bipolar disorder.

| A) Initial sample: baseline with available KSAD |  |  |  |  |
| --- | --- | --- | --- | --- |
|  | Depression | No depression | Total | Prevalence (%) |
| General | 741 | 10993 | 11734 | 6.31 |
| High-risk | 409 | 3066 | 3475 | 11.77 |
| B) MRI matched sample: depression at two-year follow-up visit |  |  |  |  |
|  | Depression | No depression | Total | Prevalence (%) |
| General | 358 | 5959 | 6317 | 5.67 |
| High-risk | 195 | 1659 | 1854 | 10.52 |
| C) Final sample: complete data at two-year follow up |  |  |  |  |
|  | Depression | No depression | Total | Prevalence (%) |
| General | 132 | 2526 | 2658 | 4.97 |
| High-risk | 59 | 566 | 625 | 9.44 |

#### 2.9 Leave-one-site-out cross-validation

Training and testing data splitting was performed using a leave-one-site-out (LOSO) cross-validation strategy (Esteban et al., 2017; Nunes et al., 2020; Sripada et al., 2021; Huang et al., 2022). This cross-validation approach takes each site separately as test data, and the steps of feature selection and model fitting via the machine learning algorithm (both described below) are applied on the remaining sites (train data). Lastly, the resulting fitted model is used to predict the outcome variable in the test data.

#### 2.10 Feature selection

Two feature selection strategies were applied to the MRI predictors. The first feature selection strategy is based on Principal Components Analysis (PCA). We extracted the first 75 principal components (following Sripada et al., 2021) from each of the multi-modal features: sMRI, DTI, task-fMRI, ROI pairwise resting-state functional connectivity. We performed an efficient PCA based on randomized algorithms (flashPCA2; Abraham & Inouye, 2014; Abraham et al., 2017). This approach can handle the eigenvalue decomposition of big datasets by decomposing it into submatrices of high probability to capture the top eigen-values and eigen-vectors. The second feature selection strategy is based on univariate analysis of variance (AOV). We tested for each predictor individually using a linear model and selected only those variables with statistically significant effects p < 0.05. However, for the pairwise functional connectivity predictors we set the threshold to p < 0.001 due to the high number of features (61776).

#### 2.11 Prediction analyses

Prediction was conducted using machine learning algorithms via the “caret” library (Kuhn, 2008; RRID:SCR_021138). Specifically, a (logistic) elastic net was applied for each type of MRI features. We select this model among other machine learning techniques because it can deal with the problem of having more predictors than observations by efficiently selecting meaningful variables (Zou and Hastie, 2005). This method has been widely used in neuroimaging (Mwangi et al., 2014; Sui et al., 2020). To deal with class imbalance (only 5-10% of cases were positive), tuning parameters were set by bootstrap and synthetic data was generated by Randomly Over Sampling Examples (ROSE; Lunardon et al., 2014). ROSE was only applied on the train data to feed balanced datasets into the elastic nets. Furthermore, due to class imbalance, prediction performance was assessed by the area under the receiver operating characteristic curve (AUROC) and the True Negative Rate (TNR) conditioned on the True Positive Rate (TPR) being over 70% (TNR|TPR>0.7).

We compared the predictive performance of each type of MRI features against other types of MRI features in a pairwise manner using a bootstrap resampling method of 10000 iterations, to account for the variation in sample size among different sites. Then, the difference in prediction performance (AUROC and TNR|TPR>0.7) was standardized based on their bootstrapped standard deviation, and its one-sided p-value was calculated to account for the polarity of the difference. Finally, all sites p-values were combined through the Fisher’s method, which multiplies minus two by the sums the p-values transformed by the natural logarithm, and this follows a chi-square distribution with degrees of freedom of two times the number of p-values (Fisher, 1992; Heard and Rubin-Delanchy, 2018). Additionally, the combined p-values were corrected for multiple comparisons using the False Discovery Rate (FDR; Benjamini and Hochberg, 1995).

## 1. 3. RESULTS

### 3.1 All MRI features predict depression onset better than chance

When assessing depression onset in the two-year follow-up, the classification performance measured by the area under the receiver operating characteristic curve (AUROC) in the whole sample (regardless of parental depression history) showed that all MRI predictors were better than random (0.5) with a 95%-confidence interval (Table 2; Figure 1; Supplementary Figure 1). The resting-state functional connectivity features extracted using univariate ANOVA (AOV) scored the highest AUROC of 0.62 (95%-CI: [0.577, 0.664]). Nevertheless, this score was not statistically significantly better than the rest of the multi-modal MRI predictors nor the extraction with the PCA feature selection. Regarding the true negative rate when setting the true positive rate to 0.7 (TNR|TPR>0.7), all scores were above the random 0.3. Similar to AUROC, rest-fMRI features with AOV were the ones with the highest score: 0.44 (95%-CI: [0.359, 0.52]).

**Table 2.** Prediction scores of depression onset at two-year follow-up for the whole sample and the subsample of the high-risk parental depression group. Numbers in parentheses represent the 95% confidence interval lower/upper bound. Scores: area under the receiver operating characteristic curve (AUROC), and true negative rate conditioned on true positive rate being over 0.7 (TNR|TPR>0.7).

| Prediction scores of depression onset at the two-year follow-up |  |  |  |  |  |
| --- | --- | --- | --- | --- | --- |
|  |  | General sample |  | High-risk |  |
| Features | Selection | AUROC | TNR\|TPR>0.7 | AUROC | TNR\|TPR>0.7 |
| sMRI | PCA | 0.58 (0.030) | 0.33 (0.081) | 0.60 (0.090) | 0.46 (0.134) |
|  | AOV | 0.59 (0.040) | 0.41 (0.079) | 0.64 (0.077) | 0.43 (0.143) |
| DTI | PCA | 0.59 (0.027) | 0.35 (0.072) | 0.62 (0.107) | 0.45 (0.162) |
|  | AOV | 0.58 (0.037) | 0.39 (0.085) | 0.66 (0.058) | 0.52 (0.098) |
| task-fMRI | PCA | 0.60 (0.046) | 0.41 (0.075) | 0.56 (0.115) | 0.43 (0.135) |
|  | AOV | 0.59 (0.044) | 0.41 (0.088) | 0.59 (0.075) | 0.42 (0.151) |
| rest-fMRI | PCA | 0.60 (0.049) | 0.40 (0.085) | 0.63 (0.065) | 0.50 (0.102) |
|  | AOV | 0.62 (0.043) | 0.44 (0.081) | 0.72 (0.065) | 0.60 (0.088) |

**Figure 1.**
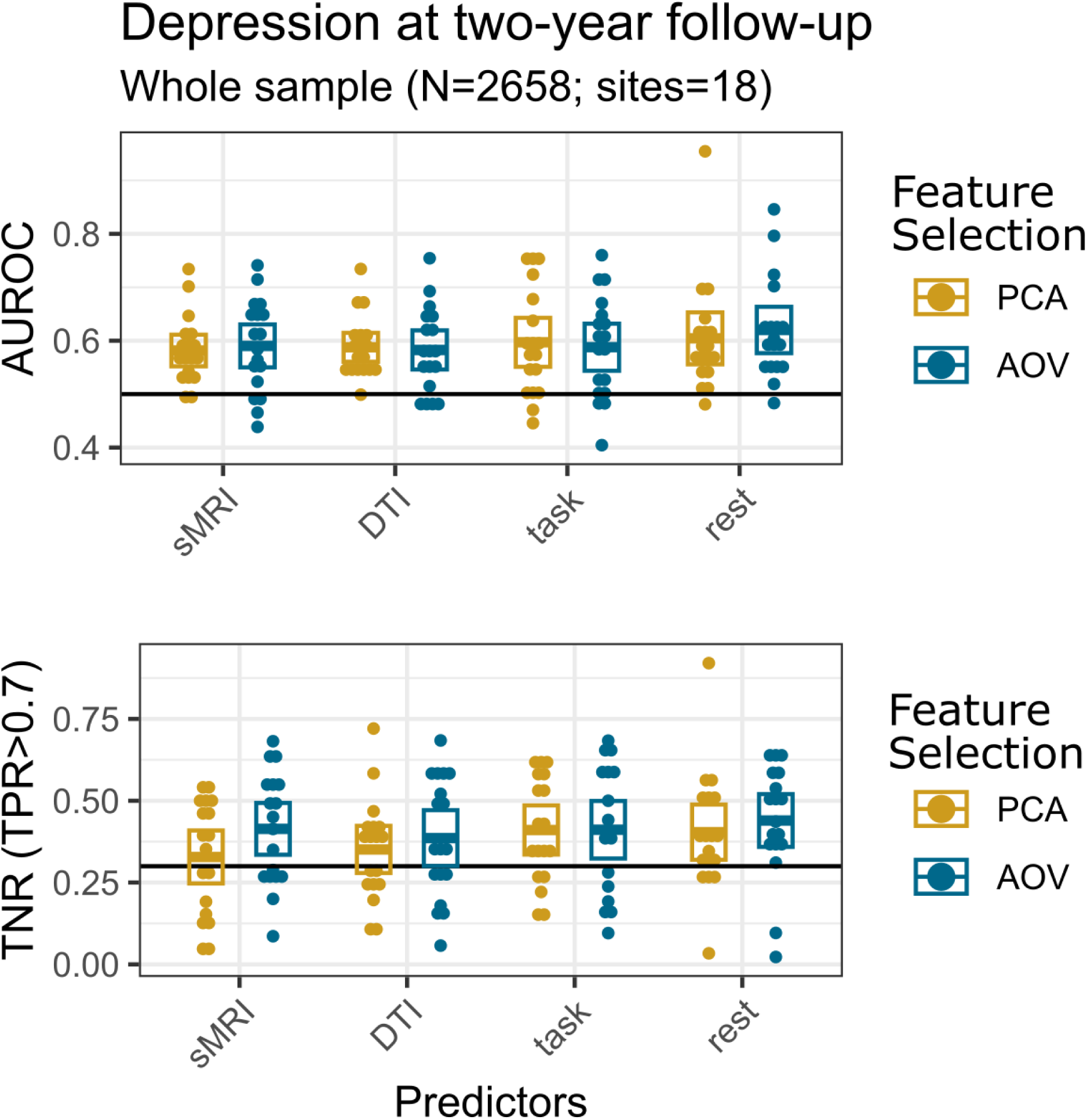
Depression onset prediction performance in the whole sample: MRI features predict depression onset better than chance. Area under the receiver operating characteristic curve (AUROC) and true negative rate (TNR) when the true positive rate (TPR) is set to higher than 0.7 scores for the prediction of depression onset at two-year follow-up, for each type of MRI predictor and feature selection method, and at each site via a leave-one-site-out (LOSO) approach. Crossbars in each boxplot indicate mean with 95%-confidence intervals. Black thick line indicates AUROC and TNR (TPR>0.7) for a random classification. Predictor abbreviations: structural MRI (sMRI), diffusion tensor imaging (DTI), task-fMRI (Task), resting-state fMRI (Rest). Feature selection abbreviations: principal component analysis (PCA), univariate ANOVA (AOV).

### 3.2 Rest-fMRI outperforms the other MRI predictors in the high-risk group

In the high-risk group with parental history of depression, the AUROC and TNR (TPR>0.7) scores for every type of features showed scores better than random (Table 1; Figure 2; Supplementary Figure 1), with rest-fMRI features via AOV showing the best performance. (AUROC: 0.72 (95%-CI: [0.651, 0.781]); TNR (TPR>0.7): 0.6 (95%-CI: [0.508, 0.683]). Furthermore, the AUROC score based on resting-state fMRI AOV features was significantly higher than DTI, task fMRI features and structural MRI features selected by PCA (Figure 3). Although it did not survive FDR correction, AUROC based on resting-state fMRI (AOV) features was marginally higher than the structural MRI (with AOV selection) (uncorrected p-value = 0.029).

**Figure 2.**
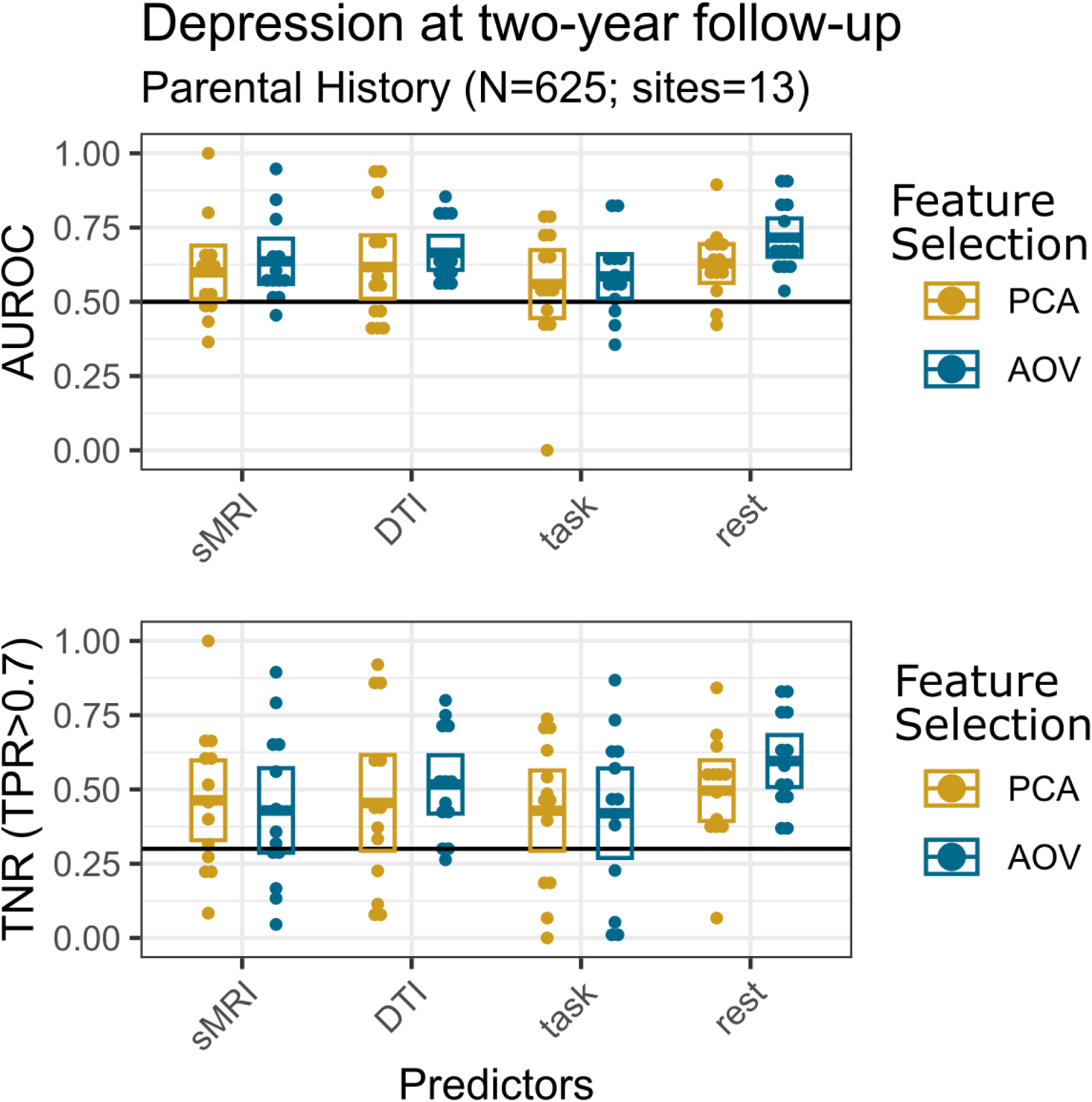
Depression onset prediction performance within the high-risk group. all MRI features predicted depression onset better than chance, with rest-fMRI features via AOV showing the best performance. Area under the receiver operating characteristic curve (AUROC) and true negative rate (TNR) when the true positive rate (TPR) is set to higher than 0.7 scores for the prediction of depression onset at two-year follow-up in the high-risk group with parental history of depression for every MRI predictor and feature selection, and at each site via a leave-one-site out (LOSO) approach. Crossbars in each boxplot indicate mean with 95%-confidence intervals. Black thick line indicates AUROC and TNR (TPR>0.7) for a random classification. Predictor abbreviations: structural MRI (sMRI), diffusion tensor imaging (DTI), task-fMRI (Task), resting-state fMRI (Rest). Feature selection abbreviations: principal component analysis (PCA), univariate ANOVA (AOV).

**Figure 3.**
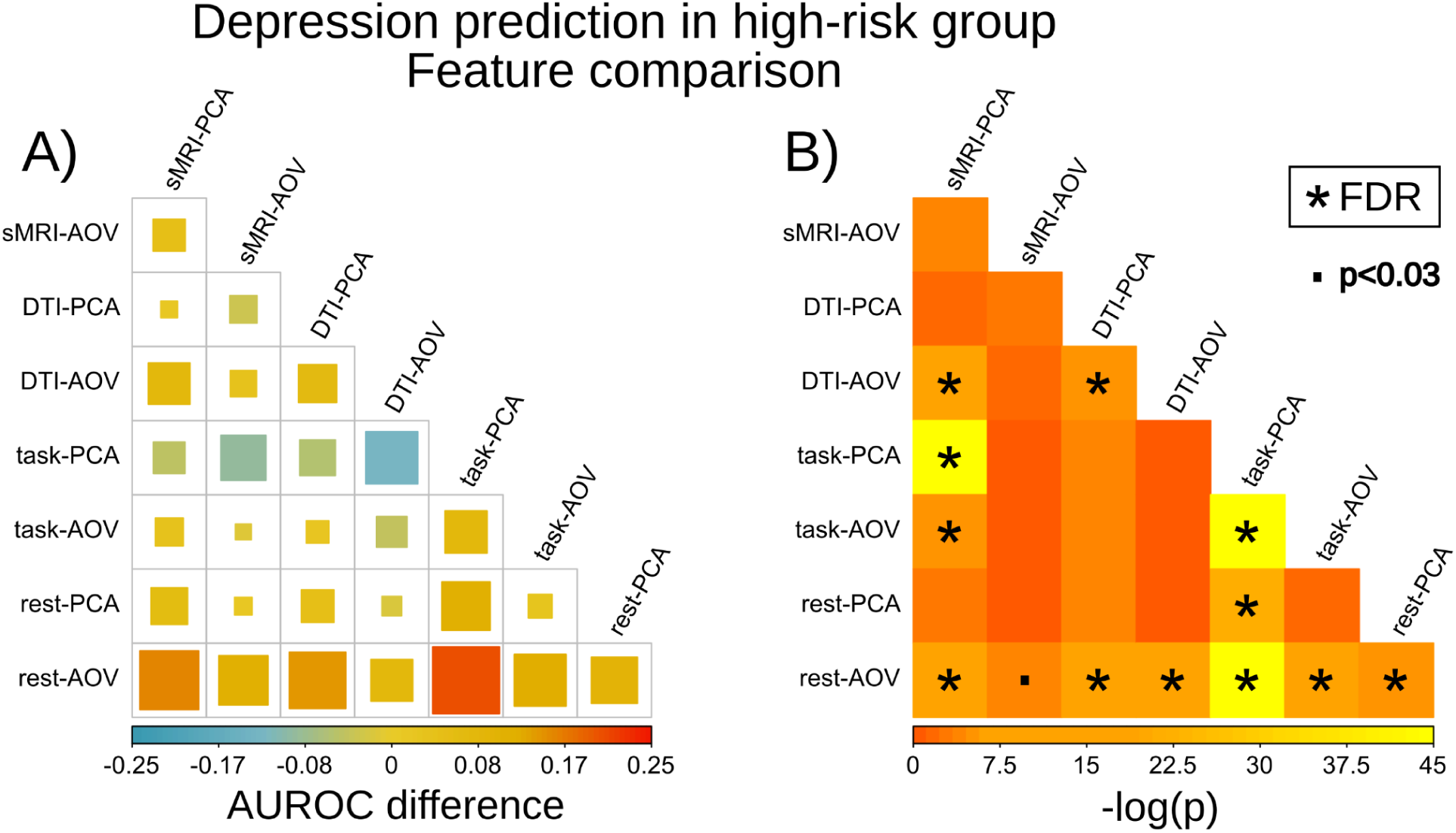
Pairwise comparisons of depression onset prediction performance of different features. rest-fMRI outperforms the other MRI predictors in the high-risk group. A: pairwise difference (weighted by site size) between the subsets of features’ area under the receiver operating characteristic curve (AUROC). The size and the color of the squares represents the raw AUROC difference of the row feature minus column feature. B: Minus natural logarithm p-value of the AUROC differences (A) of the row feature minus column feature. AUROC is based on the two-year follow-up risk of depression prediction in the subsample with parental history of depression using an elastic net classifier. P-values were calculated with the Fisher’s method that combined all sites p-values of the bootstrapped AUROC differences per site. ‘*’ denotes a significance after multiple comparison correction of FDR q < 0.05. ‘.’ denotes an uncorrected significance below 0.03.

### 3.3 Top rest-fMRI features that predicted depression onset

Given that the rest-fMRI features extracted via univariate ANOVA were the subset of features with overall higher prediction scores particularly in the high-risk group, we extracted the top features in a post-hoc analysis based on their effect size and the proportion of times they were extracted from the leave-one-site-out cross-validation (Supplementary Table 1). In the subsample with parental history of depression, those participants with higher depression risk showed an increase in their functional connectivity between the right presubiculum and the right superior frontal cortex (Area 9-46d), between the right superior parietal lobule and the left auditory cortex, between the right superior temporal sulcus and the right temporo-parieto-occipital junction, and between the left premotor (Area 6) and the right fusiform face complex. In contrast, participants with higher depression risk showed lower functional connectivity between the left operculum (OP1-SII) and the right perirhinal-entorhinal cortex (PEC), between the superior and the middle right temporal gyri, between the left sensorimotor and the left hippocampus, and within the left sensorimotor cortex.

## 1. 4. DISCUSSION

The present study aims to predict the onset of depression in a two-year span based on baseline multimodal brain imaging, using the preprocessed data from the ABCD Study sample. We also investigated the same question in a high-risk subsample of participants with parental history of depression. Our results showed that the prediction performance of brain features is significantly better than chance in both samples, with higher performance in the high-risk sample. Furthermore, when considering the high-risk subsample, the performance of the functional connectivity features extracted from the resting-state fMRI showed a good performance of AUROC = 0.72, which is significantly higher than the rest of the features from structural, diffusion, and task-based functional imaging.

A small number of recent studies have attempted the prediction of prospective onset of depression in adolescent samples using neuroimaging features. Earlier studies with small samples have shown promising prediction accuracies. For example, using relatively small samples, Foland-Ross et al. (2015) using a structural MRI feature (cortical thickness) into a 5-year follow-up sample (N=33 girls; 10-15 years old) found a 70% accuracy, where the cortical thickness of the right orbitofrontal and precentral gyri, left ACC, and bilateral insular cortex are the strongest predictors. Also using a pilot sample, Hirshfeld-Becker et al. (2019) found in a 3-4 year follow-up study of children with family history of depression (N=25; 8-14 age range at baseline) higher prediction scores based on functional connectivity compared to baseline clinical scores, on which participants developed major depressive disorder (MDD). Recent studies based on large multi-site samples have started to emerge thanks to the availability of larger open-source data. For example, Ho et al. (2022) assessed the prediction of the depression symptoms in a 1-year follow-up also in the ABCD Study (N=7995; 75/25% train/test) using phenotypical and brain (sMRI and rest-fMRI) data as features. They were able to predict slightly above a 10% of variance, with the parental history of depression among the top features, and lower functional connectivity between the right caudate and the retrosplenial-temporal network being the most relevant brain feature, although the phenotypic features were the ones with highest influence in the resulting models. Another study by Toenders et al. (2022) applied machine learning algorithms to predict depression onset in a 2- and 5-year follow-up multi-site sample (N=407/137 train/set) of an older cohort of adolescents (14-year-olds) using phenotypic and sMRI features. They found similar prediction scores (AUROC = 0.68-0.72) comparable to this study, and among the most relevant features lower surface area in the supramarginal gyrus was related to depression onset. It’s worth mentioning that when assessing our current performance with the previous multi-site approaches, our results showed decent classification scores even taking into account that we only used neuroimaging features instead of a mix of brain and phenotypic predictors from previous results (Ho et al., 2021; Toenders et al., 2022).

A few insights can be learned by considering our results along with these previous studies. First of all, prospective depression prediction when using large and multi-site samples show modest performance compared to lower samples (N<100), where the (published) scores tend to be much higher (Gao et al., 2018). This is a currently debated topic in neuroimaging, given that most published studies rely on small sample sizes which may inflate the effect size (Owens et al., 2021; Marek et al., 2022). Particularly, Winter et al. (2022) using a large (N>1800; age range: 18-65 years) and balanced sample of adults diagnosed with depression and controls found low univariate effects sizes when using multi-modal MRI predictors. Nevertheless, our results showed that brain imaging may result in good performance and medium effect sizes when applying multivariate analyses in a pediatric sample. In fact, the very tight age range may offer better predictions compared to adult populations. Given the relevance of targeting early intervention, even modest prediction scores compared to the ones in small samples may be valuable for this goal.

Lastly, when assessing several multimodal imaging, our results and previous studies showed that the functional connectivity features tend to show higher prediction scores than the other common MRI features (sMRI, DTI, and task) (Morgan et al., 2021; Ho et al., 2022; Ooi et al., 2022). This higher performance may be due to the bivariate nature of the functional connectivity, which takes into account co-activation effects or interactions between brain regions, instead of focalized ones from single brain regions such as the structural properties (sMRI and DTI) or activation patterns (task-fMRI). Even for neuropsychiatric diseases with impaired or altered behavior associated with specific tasks, resting-state data may still provide additional useful information over task-specific data for disease prognosis. This is because for disease prognosis, the individual is not yet sick, then task-specific response may still be operative for the individual. However, there may already be signs of network dysregulation in the brain functional connectome, as measured by the resting-state data. Therefore, it is likely that resting-state data are particularly useful for disease prognosis. For structural and task data the univariate approach is still the most widely used, and these single region measures were the currently available in the preprocessed ABCD repositories. However, bivariate and even more complex approaches have already been applied into structural and functional neuroimaging, such as morphometric similarity network (MSN), or task-based connectivity (Seidlitz et al, 2018; Ooi et al., 2022; Expert et al., 2019). Future research based on these network features may yield better prediction power.

Furthermore, within the group of children with parental history of depression, additional analyses of the top features from the baseline scan found several functional connectivity patterns related to the onset of depression. In addition, being able to predict better than chance in a two-year span indicates that those connections related to depression risk are already established before the start of adolescence at 9-10 years of age. Previous cross-sectional studies have shown that children and adolescents with parental history of depression have different rest-fMRI functional connectivity patterns than neurotypical samples. Particularly, lower functional connectivity between the right supramarginal gyrus (rSMG) and dorsal frontal areas were found in participants with parental history of depression (Sylvester et al., 2013; Clasen et al., 2014). In addition, Chai et al. (2016) found higher functional connectivity of the rSMG with the right amygdala, as well as within the default mode network in the group with parental history of depression. Our present results are broadly consistent with these previous findings, showing top predictive features in the lateral parietal region and DMN. These studies along with the present work demonstrate that the functional organization of the brain from those individuals with parental history of depression is different from neurotypical samples.

## 1. 5. CONCLUSION AND FUTURE DIRECTION

This work demonstrated that the onset of depression in early adolescence can be predicted from multimodal brain imaging data. Our results showed the prospective prediction performance of resting-state functional connectivity has strong predictive power for depression onset, especially within the high-risk group of children with parental history of depression. Next releases of the ABCD study, which will contain a larger set of participants who have developed depression, will provide opportunities to further explore the brain biomarker of depression in early adolescents.

## DATA AVAILABILITY

The ABCD data repository grows and changes over time. The ABCD data used in this report came from ABCD collection 3165 and the Annual Release 3.0, DOI: https://doi.org/10.15154/1519007.

## ACKNOWLEDGEMENTS

This study was partially supported by HBHL grant (XC), and Brain Canada (XC), Canada Research Chairs program (CIHR) (XC). ZGT was funded by the European Union’s NextGeneration programme and the Ministry of Universities RD 289/2021 UNI/551/2021 (Margarita Salas).

## ABCD STUDY ACKNOWLEDGEMENT

Data used in the preparation of this article were obtained from the Adolescent Brain Cognitive Development (ABCD) Study (https://abcdstudy.org), held in the NIMH Data Archive (NDA). This is a multisite, longitudinal study designed to recruit more than 10,000 children aged 9-10 and follow them over 10 years into early adulthood. The ABCD Study is supported by the National Institutes of Health and additional federal partners under award numbers U01DA041022, U01DA041028, U01DA041048, U01DA041089, U01DA041106, U01DA041117, U01DA041120, U01DA041134, U01DA041148, U01DA041156, U01DA041174, U24DA041123, U24DA041147, U01DA041093, and U01DA041025. A full list of supporters is available at https://abcdstudy.org/federal-partners.html. A listing of participating sites and a complete listing of the study investigators can be found at https://abcdstudy.org/scientists/workgroups/. ABCD consortium investigators designed and implemented the study and/or provided data but did not necessarily participate in analysis or writing of this report. This manuscript reflects the views of the authors and may not reflect the opinions or views of the NIH or ABCD consortium investigators.

