## Supplementary material for "Predicting depression risk in early adolescence via multimodal brain imaging"

Title:

Affiliations:

Supplementary methods: fMRI paradigms

fMRI sequences include four 5-minute runs of resting condition and three task-specific paradigms with two runs each. In the rest runs participants were asked to remain still with their eyes open while seeing a fixation crosshair. Task fMRI paradigms include a Monetary Incentive Delay (MID) task, a stop signal task (SST), and an emotional N-back task.

MID measures reward processing (Knutson et al., 2000; Yau et al., 2012). Each run consists of 50 pseudorandom trials (5:42 total time) that start with an incentive cue (2000 ms) of five possible types (Win 20¢, Win $5, Lose 20¢, Lose $5, or $0) followed by a jittered anticipation event (between 1500 and 4000 ms). Then, a variable target is presented (between 150 and 500 ms) and the participant will try to respond to win or avoid losing money. Next, the outcome of the trial appears as a feedback message (2000 ms minus target duration). The contrast conditions are: 1) reward vs. neutral, 2) loss vs. neutral, 3) reward vs. negative feedback, 4) loss vs. negative feedback, 5) anticipation of large reward vs. neutral, 6) anticipation of small reward vs. neutral, 7) anticipation of large reward vs. small reward, 8) anticipation of large loss vs. neutral, 9) anticipation of small loss vs. neutral, 10) anticipation of large loss vs. small loss.

SST assesses inhibition and impulse control (Logan, 1994; Whelan et al., 2012). It evaluates the withholding of a motor response (‘Go’ stimulus) when an unpredicted stop warning (‘No-Go’ stimulus) is presented subsequently. Each run has 180 trials with left or right arrows as ‘Go’ stimuli, and participants are instructed to respond in a control box the direction of the arrow as fast as possible. In 30 of those trials an upward arrow is presented as a ‘No-Go’ stimuli. Trials last 1000 ms and ‘No-Go’ stimuli vary in 50ms-interval depending on the participant performance to achieve approximately a 50/50 percent of correct and incorrect stop trial responses. The contrast conditions are: 1) correct go vs. fixation, 2) correct stop vs. correct go, 3) incorrect stop vs. correct go, 4) any stop vs. correct go, 5) correct stop vs. incorrect stop, 6) incorrect go vs. correct go, 7) incorrect go vs. incorrect stop.

The emotional N-Back task evaluates the working memory performance during emotion regulation (Cohen et al., 2016). It includes blocks of low-memory load (0-Back) where participants have to respond if the stimulus presented is the same as the one at the beginning of the block, and high-memory load (2-Back) where participants are asked to respond if the stimulus presented is the same as the one presented two trials before. Four types of stimuli were presented within the blocks: positive faces, negative faces, neutral faces, or places. Blocks include 10 trials with 4 fixation slides of 15 seconds. Trials consist of the presentation of the stimulus for 2 seconds followed by a fixation cross for 500 ms. A total of 160 trials were presented, 80 for each memory-load condition and 40 for each type of stimulus. The contrast conditions are: 1) 0-Back condition, 2) 2-Back condition, 3) Place condition, 4) emotion condition, 5) 2-Back vs. 0-Back, 6) face vs. place, 7) emotion vs. neutral face, 8) negative face vs. neutral face, 9) positive face vs. neutral face.

Supplementary Figures

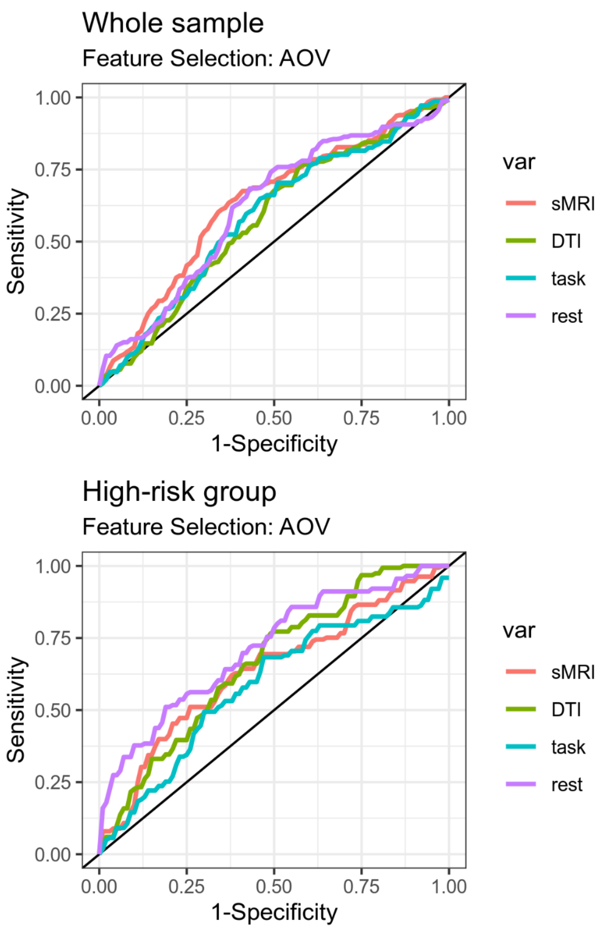


Supplementary Figure 1. Averaged Receiver Operating Characteristic (ROC) curves for the general sample (top) and high-risk group with parental history of depression (bottom), by feature selection approach: univariate ANOVA (AOV) (left) and principal component analysis (PCA) (right), for each MRI predictors: structural MRI (sMRI), diffusion tensor imaging (DTI), task-fMRI (task), and resting-state fMRI (rest).

Supplementary table 1. top rest-fMRI features that predicted depression onset. rest-fMRI features with higher effect (|Cohen's d| > 0.5) on depression onset in the high-risk group. Features correspond to functional connectivity edges extracted through the Leave-One-Site-Out (LOSO) cross-validation. ROI 1 and ROI 2 account for the regions of interest of the corresponding edge, LOSO % accounts for the percentage of sites in which that feature was selected as predictors after the univariate ANOVA feature selection. R or L indicates the right or left hemisphere.

| ROI 1 | ROI 2 | LOSO % | Cohen's d |
|---|---|---|---|
| Area OP1-SII (L) | Perirhinal Ectorhinal Cortex (R) | 100 | -0.60 |
| PreSubiculum (R) | Area 9-46d (R) | 92.3 | 0.54 |
| Auditory 4 Complex (L) | Area 7PC (R) | 92.3 | 0.53 |
| Area 6mp (R) | Area 7PC (R) | 84.6 | -0.54 |
| Area STSv posterior (R) | TemporoParietoOccipital Junction 2 (R) | 84.6 | 0.51 |
| Area STSv posterior (R) | Area TE2 anterior (R) | 84.6 | -0.50 |
| Area 5m (L) | Inferior 6-8 Transitional Area (R) | 84.6 | -0.52 |
| Auditory 4 Complex (L) | Lateral Area 7A (R) | 84.6 | 0.51 |
| Primary Sensory Cortex (L) | Hippocampus (L) | 76.9 | -0.51 |
